## SupplementalMethodsResultsFigures for "Social and environmental predictors of gut microbiome age in wild baboons"

**Affiliations**


**File contents**


Supplementary Text
Supplementary Figures
- Figure S1
- Figure S2
- Figure S3
- Figure S4
- Figure S5
- Figure S6
- Figure S7

### Supplementary Text

<u>Creating and assessing age-predictive machine learning models</u>

*Introduction to the approaches.* To create our final microbiome aging clock, we tested three supervised machine learning algorithms: elastic net regression, Random Forest regression, and Gaussian process regression (*107*, *109*, *110*). Below we summarize the strengths and weaknesses of each machine learning algorithm.

Elastic net regression is a regression algorithm that produces a linear model. It improves upon the predictions from simple linear regressions by incorporating coefficient penalties from the L1 regularization (LASSO regression) and L2 regularization (ridge regression) (*109*). Elastic net regression is infrequently used in microbiome studies but has produced promising results in epigenetic aging clocks due to its flexibility in choosing which features to keep and which to remove (*40–43*, *45*). However, elastic net regressions produce linear relationships between the input chronological age and the predicted age, which may not accurately affect the true relationship between chronological age and the microbiome.

Random Forest regression is an ensemble learning method that creates a number of parallel decision trees, each producing its own prediction (*110*). The prediction is then averaged among all trees to create the final estimate. A key advantage of Random Forest regression over elastic net regression is that it does not assume a linear relationship between the predicted estimate and the input chronological age, but the model may be biased by correlated features. Random Forest regression is commonly used in microbiome research, including other microbiome clocks (*36*, *111–113*).

Gaussian process regression is a nonparametric, Bayesian approach that infers a probability distribution over all the potential functions that fit the data (*107*). Like random forest, Gaussian process regressions do not assume a linear relationship between chronological age and predicted age but has the additional advantage of kernel customization. As such, Gaussian process regressions may be able to better handle heteroskedasticity in the data (an issue in our clock; see below). As an increase in chronological age is often associated with the breakdown of physiological processes (e.g. aging), heteroskedasticity in microbial age estimates may indicate a breakdown of the host's processes that regulate the gut microbiome.

*Methods and optimization of machine learning algorithms.* Prior to running each algorithm, all features were center log ratio transformed within sample. We then chose a ratio of training to test dataset. To do this, we first compared the model fit of different ratios of training to test sets. These included the following training:test splits: 50:50, 60:40, 75:25, 80:20, and 90:10. We found that an 80:20 data split provided the best balance between model performance and the risk of overfitting.

In order to calculate a microbial age estimate for every sample and estimate generalization error, we used a nested cross-validation framework. Each of the three algorithms has its own internal cross-validation where a subset of the training data is held apart and used to internally validate the model. We added an additional, external layer of cross-validation with our 80:20 training:test data split. We classified samples into five different test sets where individual was as evenly represented as possible in all training and test sets. As the number of samples varied between individuals, we randomly assigned each sample a test set without replacement if an individual's sample count was less than five, or with replacement if an individual's sample count was greater than five. For each model run, four of the test datasets were treated altogether as training data and the fifth set was the validation test set.

Elastic net regressions were run in R using function cv.glmnet() from package glmnet (*114*). The two main parameters for this model are $\lambda$, which is the penalty from the LASSO regression that penalizes extra predictors by shrinking coefficients to zero, and $\alpha$, the parameter that balances between minimizing between the residual sum of squares and minimizing the magnitude of the coefficients. cv.glmnet() automatically fits 100 values of $\lambda$ by default and names the $\lambda$ that produces the minimum cross-validated error "lambda.min". We used lambda.min as our value of $\lambda$. For $\alpha$, we manually ran the model with 200 values of alpha (from 0 to 1 in increasing increments of 0.005) and picked a value of alpha that would minimize the mean absolute error and maximize the adjusted $R^2$.

Random Forest regressions were conducted in Python 3 using scikit-learn (*105*). The main parameter was the number of decision trees being used, which defaults to 100. Too many trees could result in overfitting so in order to minimize overfitting and optimize $R^2$, we ran a series of Random Forest regressions with different numbers of trees: we increased the number of trees in increments of 50, stopping at 400 because of minimal changes in $R^2$ relative to 200 trees.

Gaussian process regressions were also conducted in Python 3 using scikit-learn (*105*, *106*). In both the non-heteroskedastic-kernel model and heteroskedastic-kernel model, the main parameters we used to modify the kernel function included the scale and bounds. These parameters moderate the level of overfitting in the algorithm: the scale parameter specifies a starting point for which the algorithm optimizes within the confines of the bounds parameters. As with the other models, we incrementally changed both the scale parameter within a wide range of bounds and checked the output model's $R^2$ and median error. Our final model retained a wide range of bounds (1 to 100) and set the scale parameter to the median euclidian distance of the dataset as calculated in R using function vegdist() from R package vegan (*102*).

Due to the heteroskedasticity exhibited by the models above (**Figure S7**), we modified the Gaussian process regression's kernel function further to account for the variance within the dataset. Specifically, we multiplied the variance in the training data by the radial basis function, which distributed the higher variance in later life more evenly across lifespan.

*Comparison of machine learning algorithms.* To assess model accuracy, we used the predicted age estimates from all 5 runs of the nested cross-validation procedure to assess model fit and accuracy. As in Horvath (2013), we regressed the sample's predicted microbial age ($age_m$) against the host's known chronological age ($age_c$) and calculated: (1) the $R^2$ between $age_c$ and $age_m$; (2) the Pearson's correlation coefficient between $age_c$ and $age_m$; and (3) the median error as the median absolute difference between $age_c$ and $age_m$ (**Table S13** and **Figure S3**). Across all algorithms, we observed that males always aged faster than females, which is consistent with well-known patterns of sex-specific senescence in humans and other primates (*84*) (**Figure S3**). The Gaussian process regression with the heteroskedastic kernel was the best model for every metric assessed - it maximized $R^2$ and Pearson's R to 0.488 and 0.698 (respectively) while minimizing median error. It also was the only model with which we were able to alleviate any heteroskedasticity.

**Supplementary Figures**

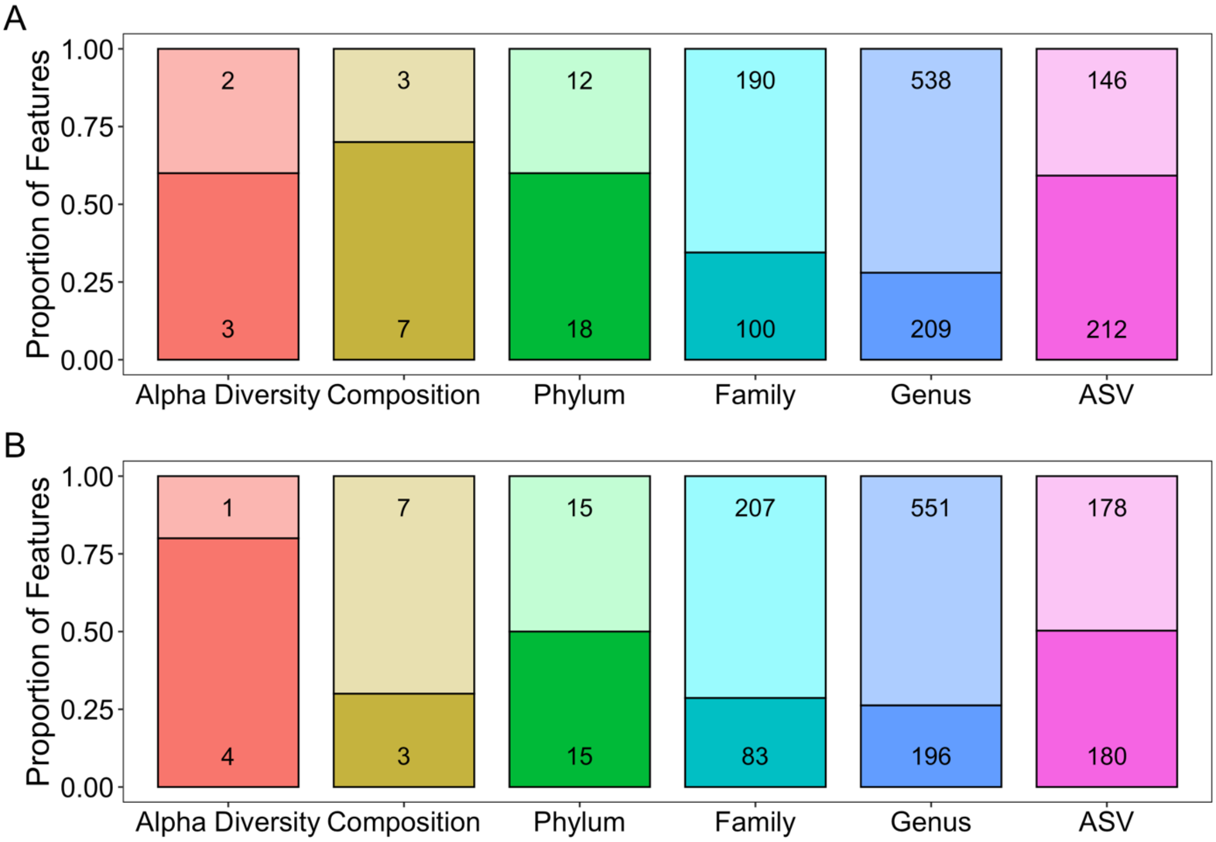


**Figure S1**. **The number and proportion of each type of feature that was significantly associated with age.** In the two panels, age was modeled as (A) a linear term, or (B) a quadratic term in a linear mixed model (FDR threshold=0.05; darker colors represent the proportion of statistically significant features). All features were modeled using Gaussian error distributions. As described in the Results, feature types included (i) five metrics of alpha diversity, the top 10 principal components of Bray-Curtis dissimilarity (collectively labeled 'Composition' in the gold bar), the abundances of each microbial phylum (n=30), family (n=290), and genus (n=747), and ASVs detected in >25% of samples (n=358).

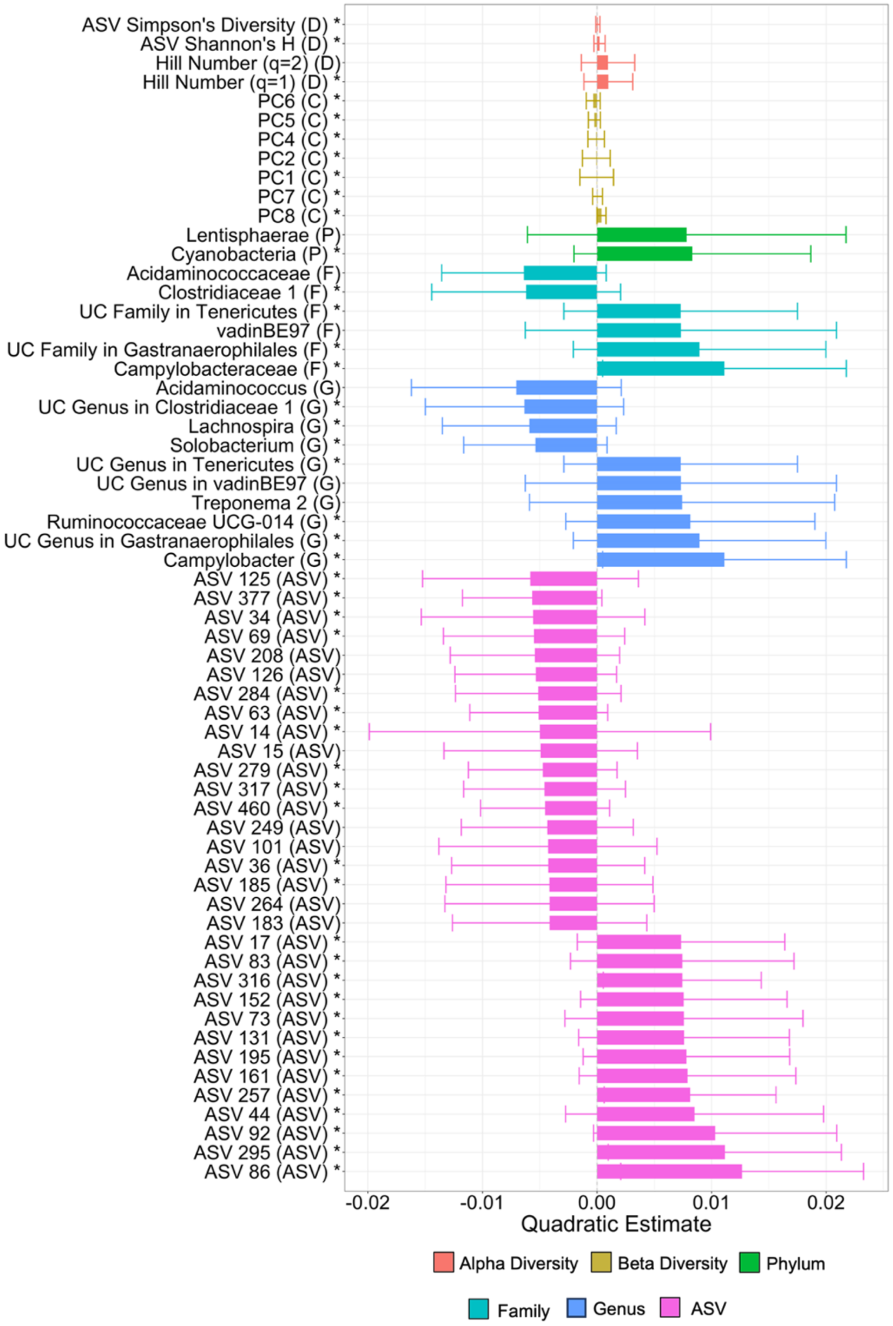


**Figure S2**. **Taxa with the strongest quadratic associations with age**. Plot shows the size of the quadratic estimate for each taxon that had a significant association with age. Bars are colored by the type of feature (see legend) and indicated by the letter in parentheses, with D indicating a diversity metric, C a compositional metric (i.e., a principal component of Bray-Curtis similarity), P for phylum, F for family, G for genus, and ASV for an ASV. To make our quadratic terms more interpretable, we centered our age estimates on zero by subtracting the mean of age from each age value. Specifically, when a quadratic term is negative, the curve is concave, whereas when the term is positive, the curve is convex. UC is short for uncharacterized. Features that also had a significant linear age term are indicated by a *.

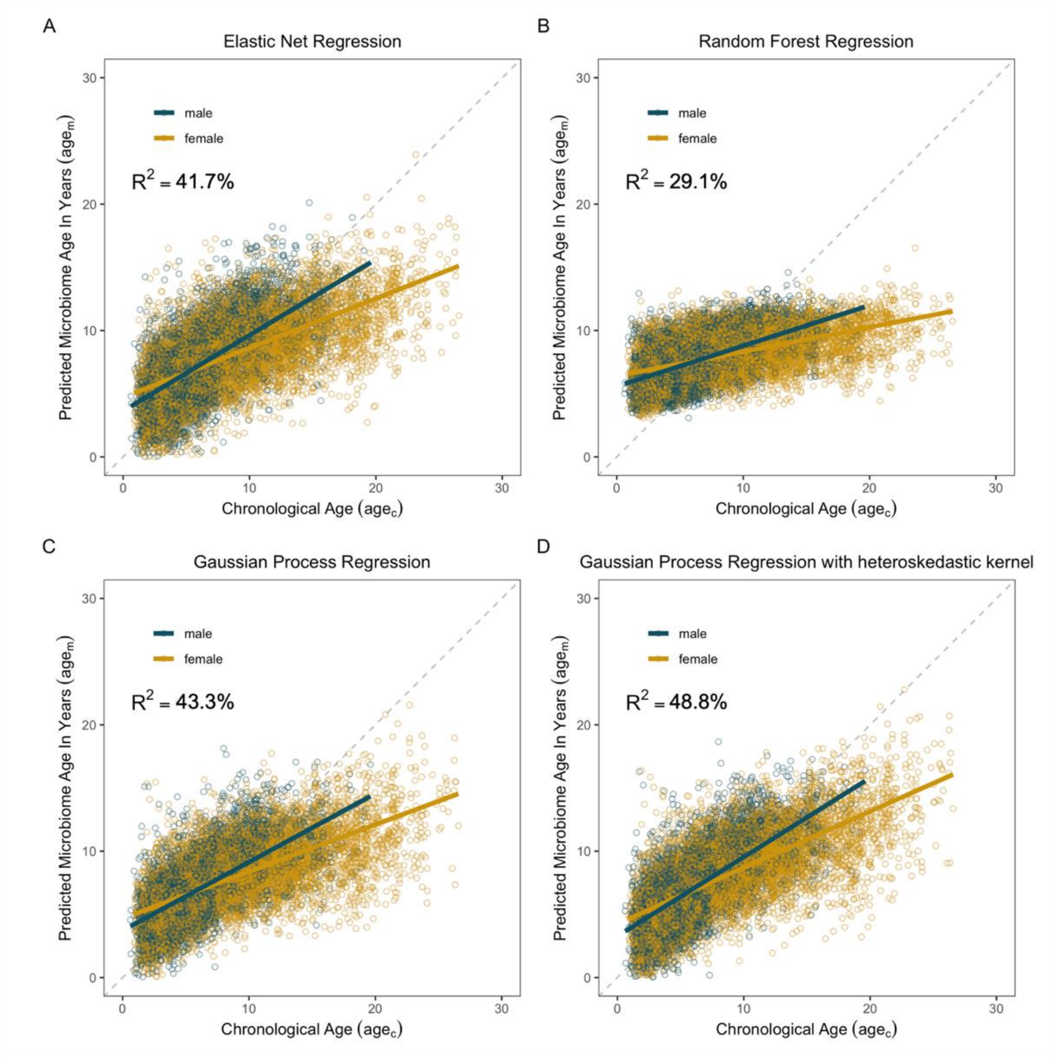


**Figure S3**. **Microbiome clocks from an ensemble of machine learning algorithms**. Each plot shows predicted microbiome age ($age_m$) relative to the true, chronological age ($age_c$) of the baboon at the time of sample collection. (A) shows age predictions from an elastic net regression, and (B) depicts age predictions from Random Forest regression. Plots (C) and (D) show age predictions from Gaussian process regression without (C) and with (D) a kernel to account for heteroskedasticity. The most accurate age predictions (i.e., the model with the highest $R^2$ value) were produced by a Gaussian process regression model with a kernel customized to account for heteroscedasticity (D). On each plot, points are colored by host sex; yellow indicates samples from females; blue indicates samples from males. Grey dashed lines indicate a 1-to-1 relationship between $age_c$ and $age_m$.

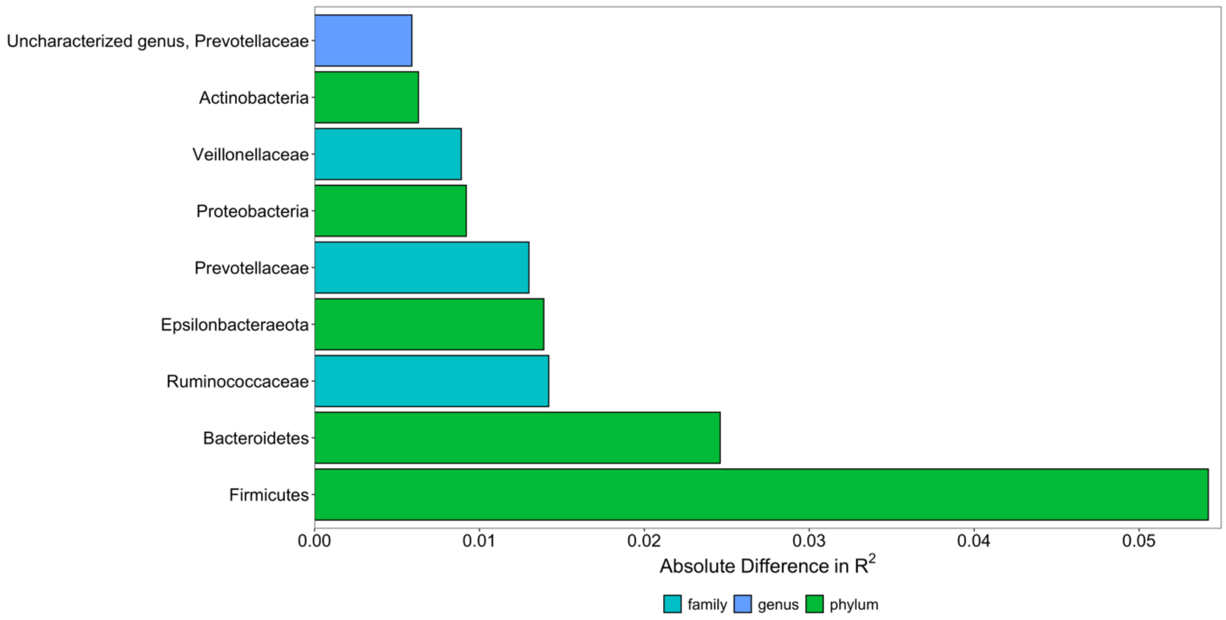


**Figure S4**. **Some microbiome features that had a substantial effect on age predictions**. Plot show the nine microbiome features that, when removed from the microbiome clock, reduced $R^2$ for the relationship between $age_m$ and $age_c$ by more than half a percent. These nine were the only ones out of the 1,081 non-ASV features we examined that exhibited this effect. The magnitude of the difference in $R^2$ with and without that feature is shown on the x-axis.

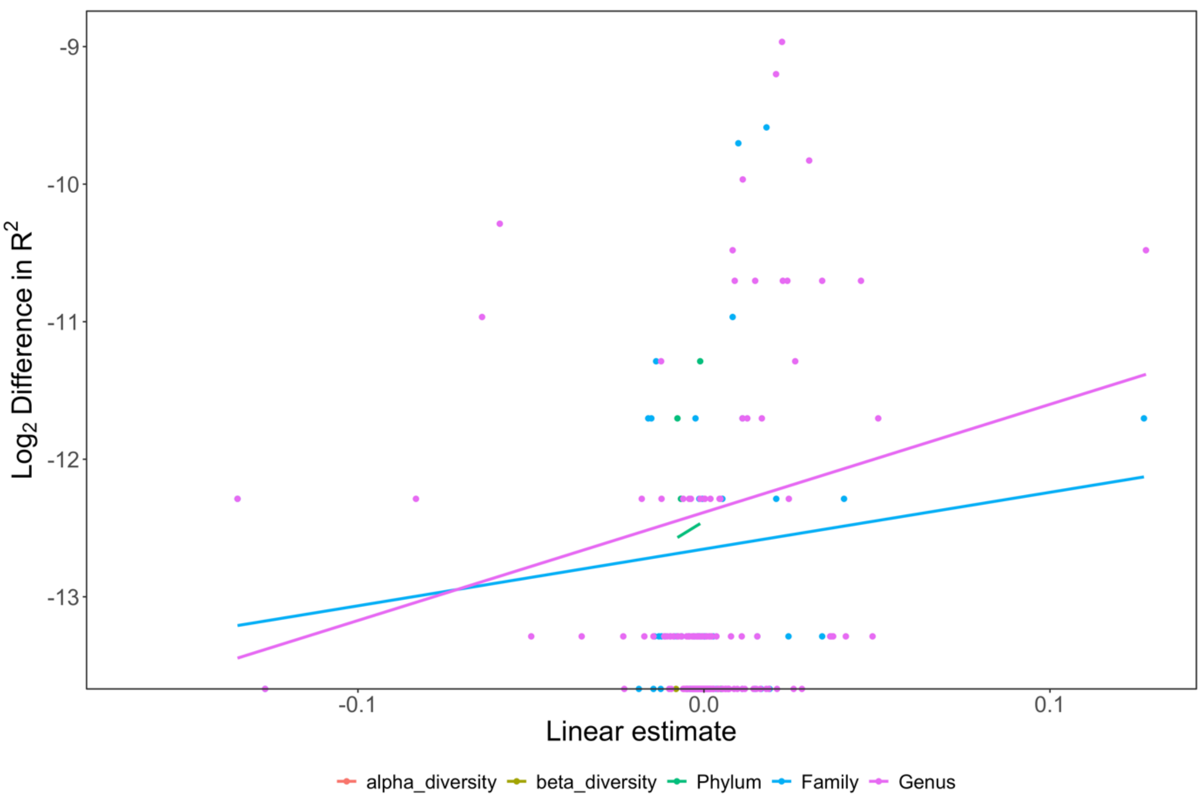


**Figure S5**. **Age associated traits had stronger effects on clock performance**. We found a weak, positive relationship between the change in $R^2$ from removing a given feature from the microbiome clock and the feature's linear relationship with age (Pearson's correlation: 0.06; p=0.023). The x-axis shows the linear coefficient for age produced when age is regressed on the microbiome feature of interest (**Table S1**). The y-axis shows the log transformed (base 2) of the difference in $R^2$ from the correlation between $age_m$ and $age_c$ without the feature compared to the full microbiome clock, including the missing feature. Points represent all non-ASV features, and lines represent the trends for that feature type.

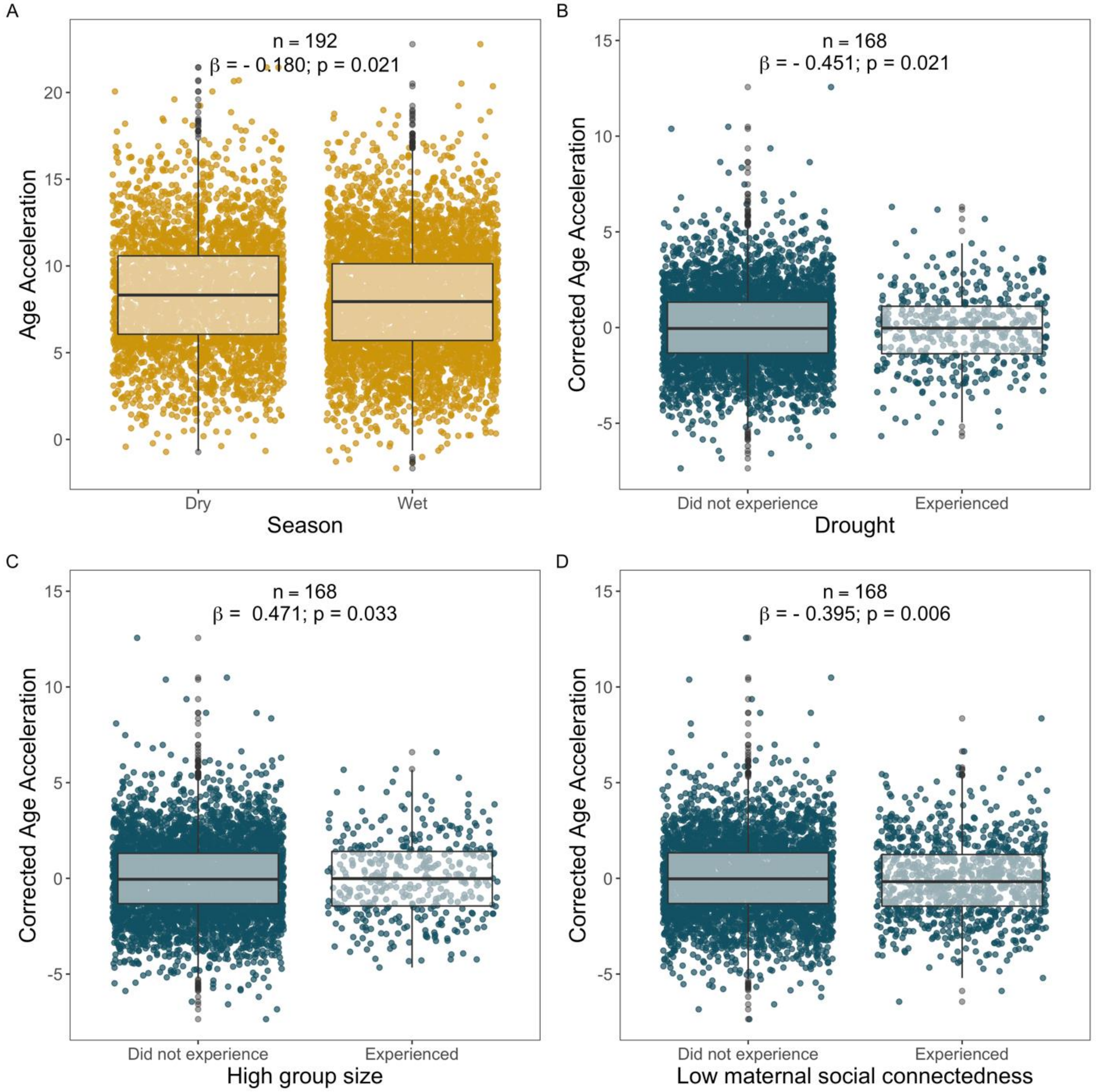


**Figure S6**. **Statistically significant socio-environmental predictors of corrected Δage not shown in Figure 4 in the main text**. Each point represents a sample: yellow points show samples from females, and blue points show samples from males. (A) Season was a weak but significant predictor of lifetime Δage in females (**Table S6A and C**). Panels (B-D) show that males who experienced (B) drought, (C) high group size at birth, or (D) low maternal social connectedness at birth exhibited variation in Δage, but in different directions: the experience of early-life drought and maternal social isolation predicted young-for-age gut microbiomes in males, while being born in a large group predicted old-for-age microbiomes (see Results for details). Corrected Δage represents the residuals of the relationship between $age_m$ and $age_c$ correcting for chronological age, season, monthly temperature, monthly rainfall, social group at the time of collection, and hydrological year (**Tables S6B, D, and E**).

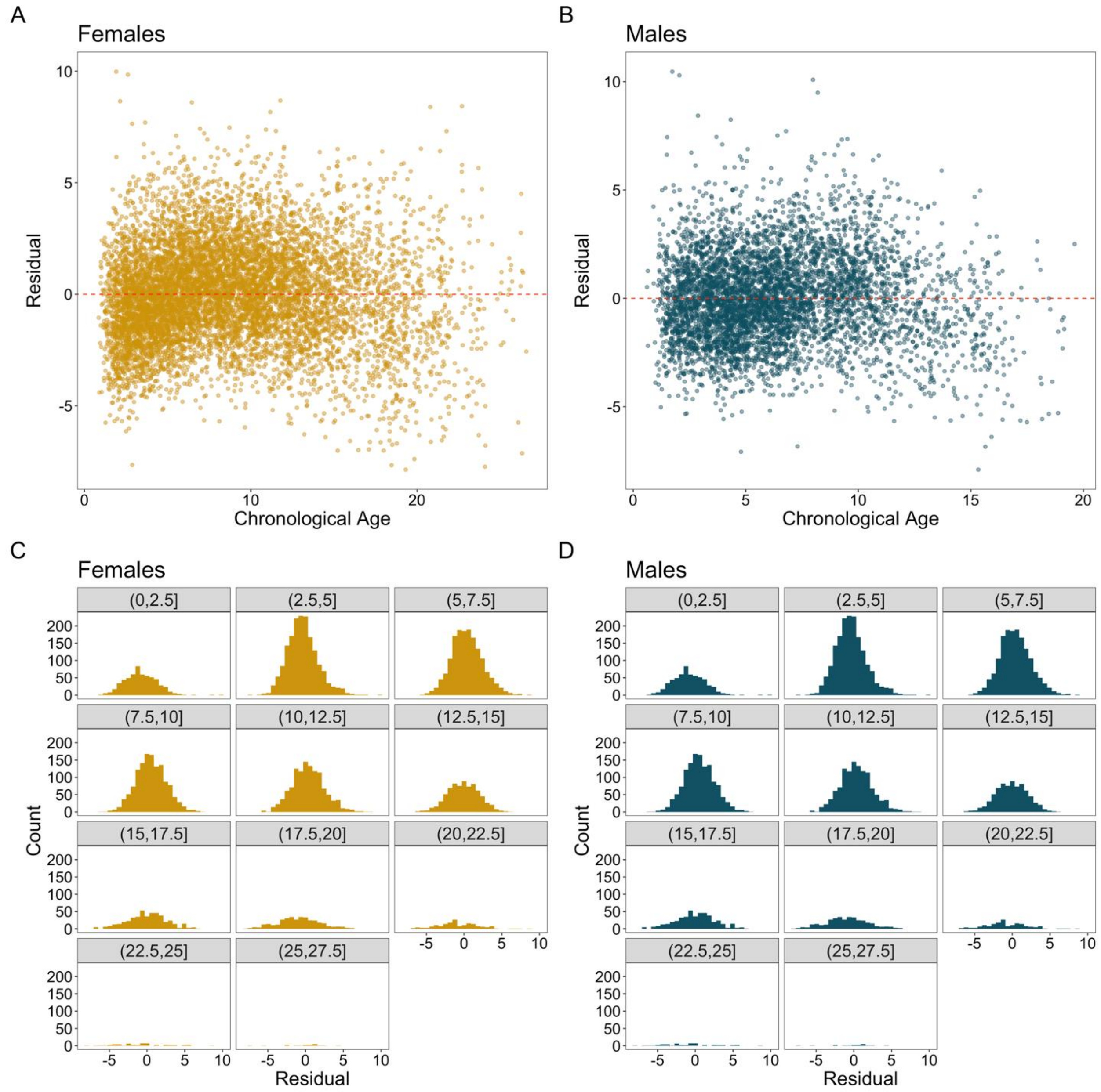


**Figure S7**. **Variance in residuals across lifespans in the Gaussian process regression prior to correction**. Plots show chronological age relative to the residuals of the $age_m$ produced by a Gaussian process regression with a radial basis function kernel. Females are in yellow, and males are in blue. (A) and (B) show a scatter plot of $age_c$ and the residuals of $age_m$. The spread of the residuals is wider for samples collected at older ages. (C) and (D) show the distributions of the residuals for different age subsets. The distribution flattens around 12.5 in females and 10 in males.

202
